# Targeted intracellular delivery of molecular cargo to hypoxic human breast cancer stem cells

**DOI:** 10.1101/2024.01.12.575071

**Authors:** Ashley V Makela, Anthony Tundo, Huiping Liu, Doug Schneider, Terry Hermiston, Pavlo Khodakivskyi, Elena Goun, Christopher H Contag

## Abstract

Cancer stem cells (CSCs) drive tumorigenesis, are responsible for metastasis, and resist conventional therapies thus posing significant treatment challenges. CSCs reside in hypoxic tumor regions and therefore, effective therapies must target CSCs within this specific microenvironment. CSCs are characterized by limited distinguishable features, however, surface displayed phosphatidylserine (PS) appears to be characteristic of stem cells and offers a potential target. GlaS, a truncated coagulation protein that is internalized after binding PS, was investigated for intracellular delivery of molecular payloads to CSCs. Intracellular delivery via GlaS was enhanced in patient-derived CD44+ mammary CSCs under hypoxic conditions relative to physoxia or hyperoxia. *In vivo*, GlaS successfully targeted hypoxic tumor regions, and functional delivery of molecular cargo was confirmed using luciferin conjugated to GlaS via a disulfide linkage (GlaS-SS-luc), which releases luciferin upon intracellular glutathione reduction. Bioluminescence imaging demonstrated effective GlaS-mediated delivery of luciferin, a model drug, to CSCs in culture and *in vivo*. These findings offer the promise of directed delivery of therapeutic agents to intracellular targets in CSCs.

## Introduction

Although there have been significant advancements in breast cancer therapy, the challenge of effectively eliminating breast cancer stem cells (CSC) as drivers of disease progression remains. CSCs are functionally defined as cancer cells with self-renewal and tumorigenic properties, that often are quiescent, undergo differentiation, and are metastatic^1–4^. CSCs tend to localize in hard-to-reach hypoxic regions of the tumor; all of these characteristics render CSC resistant to traditional therapies (*i*.*e*., chemotherapy, radiation therapy, and certain molecular targeted drugs). Residual disease is likely due to conventional therapies failing to eliminate CSC concurrently with the bulk of the tumor, allowing the CSC to persist and subsequently relapse, differentiate and metastasize. The necessity to eliminate CSC is paramount for effective treatment, and methods for targeting the CSC population are essential. However, CSCs exhibit few characteristics that allow them to be distinguished and targeted. However, there are relevant CSC characteristics that we aimed to target in this study, including outer leaflet phosphatidylserine (PS) exposure on the cell surface^5^, CD44 expression^6^, and a tendency to reside in the hypoxic tumor microenvironment (TME)^6^.

Hypoxia is common within solid tumors, and in the context of breast tumors, can be as low as 1% O2 (severe hypoxia), compared to physoxia (oxygen levels found in normal tissues) at 3-7% O2^7^. In contrast, cells in culture are typically propagated in hyperoxia at 21% O_2_. Further, the stem cell niche operates under reduced oxygen conditions of 2-6% O_2_^8^, and may be involved in supporting stem cell characteristics. Hypoxia inducible factors (HIFs; HIF1α and HIF2α) are transcription factors that are expressed during exposure to decreased O_2_, and their activity regulates the expression of many stem cell markers^9–11^. In addition, hypoxia combined with radiotherapy seems to enhance the stem-like properties of cells^12^. In the context of this study, we used three experimental oxygen levels; 7% for physoxia, 4% for hypoxia (mild) and 21% for hyperoxia. Typically, studies of CSC utilize cell culture conditions at room oxygen levels (21%) and control oxidative stress by adding antioxidants to the media^13^. However, no cells in internal organs, with the exception of cells lining the alveoli in the lungs, grow in 21% oxygen, and in the tumor microenvironment the oxygen levels are well below 21%. The discrepancies in oxygen concentrations in culture verses in organs and tissue, can potentially lead to erroneous results when investigating cellular characteristics and responses to investigative therapeutics^14,15^. Consequently, studying CSC under conditions that approximate their niche in the TME, specifically with regard to oxygen levels, is critical to evaluating innovative CSC targeting therapeutics.

PS is a negatively charged glycerophospholipid present in all eukaryotic cell membranes. It is generally inaccessible as a target since it remains a component of the inner leaflet through the action of ATP-dependent aminophospholipid flippase in most physiologically healthy cells^16^. The externalization of PS is observed under some biological processes including necrosis and apoptosis^17^, blood coagulation^18^, disease^19,20^ and on stem cells^21^. Dying cells lack ATP, and thus PS is exposed on their surface subsequently leading to them being eliminated by nonspecific phagocytosis by immune cells. PS externalization appears to signal the immune system not to mount an immune response to antigens on these cells since they are indicators of “self” thus avoiding autoimmune responses. As such, tumor cells that have increased PS on their outer membrane^19,20^ and/or display elevated levels of CD47^22^ on their surface evade immune destruction. Thus, PS externalized on cancer cells is an immunosuppressive mechanism, facilitating tumor growth and enabling metastasis^23–25^. PS-binding proteins (*i*.*e*., Annexin V; AnnV)^26,27^ and mAbs (*i*.*e*., Bavituxumab; Bavi)^28^ are being studied as a means of targeting PS-expressing cancer cells. Bavi works to reverse the immunosuppressive actions of the externalized PS^29^, acting as an immunomodulatory therapy; although these proteins bind to PS on the cell surface, they are not internalized, limiting their use to the delivery of surface-active agents. In contrast, we have found that a truncated protein S comprised of the PS-binding γ-carboxyglutamate (Gla) domain and four epidermal growth factor (EGF) domains (collectively referred to as GlaS), binds PS on the outer leaflet and is internalized after binding^30^ enabling delivery of payloads to the cytoplasm of cells with externalized PS, which would include those in the hypoxic regions of tumors.

Drug delivery mechanisms can be categorized into passive and active targeting. Passive targeting, driven by the enhanced permeability and retention (EPR) effect, leads to drug accumulation in the tumor; however, this approach is generally unreliable and often results in low drug delivery efficiency^31–33^. In contrast, adding a targeting moiety significantly enhances therapeutic efficacy compared to passive targeting^34^. Active targeting leverages the expression of specific markers in the TME as docking sites for mAbs, proteins, or peptides. Antibody-drug conjugates (ADCs), which enable intracellular drug release after internalization, have demonstrated improved therapeutic efficacy and safety both *in vitro* and *in vivo*^*35,36*^. Although these mechanisms have increased therapeutic efficacy, there are still problems such as off-target delivery and heterogeneity of the biomarkers, leading to inconsistent therapeutic effect. A universal target, for example PS, for translation across the plasma membranes of different cancer types would have tremendous utility in cancer therapy since PS is unlikely to be down regulated in response to the selective pressure of a therapeutic. Levels of externalized PS have been evaluated in a variety of cancers using radiolabeled PS binding proteins; in these studies PS signals are weak prior to initiation of therapy and increase with cell death^37–39^.

Validation of new PS targeting moieties as intracellular delivery tools can first be achieved through imaging. An imaging reporter, such as the small molecule D-luciferin, can serve as a model drug mimetic, allowing for the visualization of intracellular delivery via bioluminescence imaging (BLI); extracellular luciferin will not lead to a bioluminescent signal. In this study, the CSCs were engineered to express the luciferase enzyme, enabling their identification using BLI when the substrate D-luciferin is internalized and co-factors such as ATP and O2 are present. D-luciferin can be conjugated to GlaS via a disulfide linkage (GlaS-SS-luc), which leads to release of luciferin from GlaS when the conjugate is internalized and exposed to intracellular concentrations of glutathione. Release of luciferin enables bioluminescence, resulting in light emission constituting a molecular light switch that reveals internal delivery. The presence of light following the delivery of GlaS-SS-luc will validate: 1) PS binding of GlaS, 2) internalization of GlaS and its cargo (luciferin), 3) release of the cargo, and 4) functional activity of the cargo.

This study examines the effects of oxygen conditions on the targeted binding and internalization of the protein, GlaS, in CSC derived from triple-negative human breast cancer patients as marked by CD44 protein expression on lineage negative tumor cells. We show that there is increased PS externalization on CSC when cultured in hypoxic conditions (4%) relative to physoxia (7%) and hyperoxia (21%), and that there are more CD44+GlaS+ cells by flow cytometry in hypoxia. Further, a fraction of small-sized CD44+ cells have increased levels of GlaS staining, suggesting that the GlaS may more effectively target a subset of more highly stem-like, quiescent and tumorigenic CSC. Using imaging, we validated internalization, release and functionality of a GlaS-conjugated small molecule (luciferin), with bioluminescence output relating to GlaS-SS-luc delivery into the CSC in culture and *in vivo*.

## Results

### Frequency of CD44+ cells and CD44+GlaS+ cells at three oxygen concentrations

Breast cancer patient-derived CD44+ CSCs cannot be maintained in culture and were therefore propagated in NOD-scid gamma (NSG) immunocompromised mice as human patient-derived xenografts (PDXs). Although there are several markers of CSC, CD44+ cells, under these conditions, are the CSC population. Five PDX lines -M1L2T, M2L2T, M3L2G, CTC205 (L2T) and CTC092 (L2T) - were evaluated. The M1 was derived from pleural effusion metastasis, M2 and M3 of human breast tumors^40^ and CTC205 and CTC092 were derived from circulating tumor cells of patients with breast cancer^41,42^. The L2T refers to labeling with the Luc2-tdTomato reporter, and L2G to labeling with the Luc2-eGFP^40^. The PDX tumors had variable growth rates in mice, and the lines M1L2T and CTC205 grew the fastest, followed by M2L2T and M3L2G, and CTC092, respectively. After the cells were harvested from dissociated tumor tissue they were placed in hypoxic (4%), physoxic (7%) or hyperoxic (21%) conditions overnight. Following overnight culture, cells were stained and examined by flow cytometry. The percent of CD44+ cells from each PDX tumor varied slightly, although none had significant differences in the percent of CD44+ cells under different oxygen conditions; there was an average of 62% CD44+ cells in hypoxic, 64% in physoxic and 60% in hyperoxic conditions (F_2,36_=1.32, *p*=.28) from live cells (**Fig. 1a**,**d**). Within populations there were no differences in the frequency of CD44+ cells (**Supplementary Fig. 1**) between the oxygen conditions in M1L2T (F_2,6_=4.89, *p*=.055), M2L2T (F_2,5_=2.13, *p*=.21), M3L2G (F_2,6_=2.26, *p*=.19) and CTC092 (F_2,1_=56.2, *p*=.09) cells. However, there were differences in the frequency of CD44+ cells in populations held at different oxygen conditions for the CTC205 (F_2,6_=11.96, *p*=.008) cells, with lower percentages of CD44+ cells in the hyperoxic condition vs the hypoxic (*p*=.049) and physoxic (*p*=.007) conditions. Despite negligible differences in the frequency of CD44+ cells in the three oxygen concentrations, the frequency of CD44+GlaS+ cells did vary with oxygen concentration.

**Figure 1.**
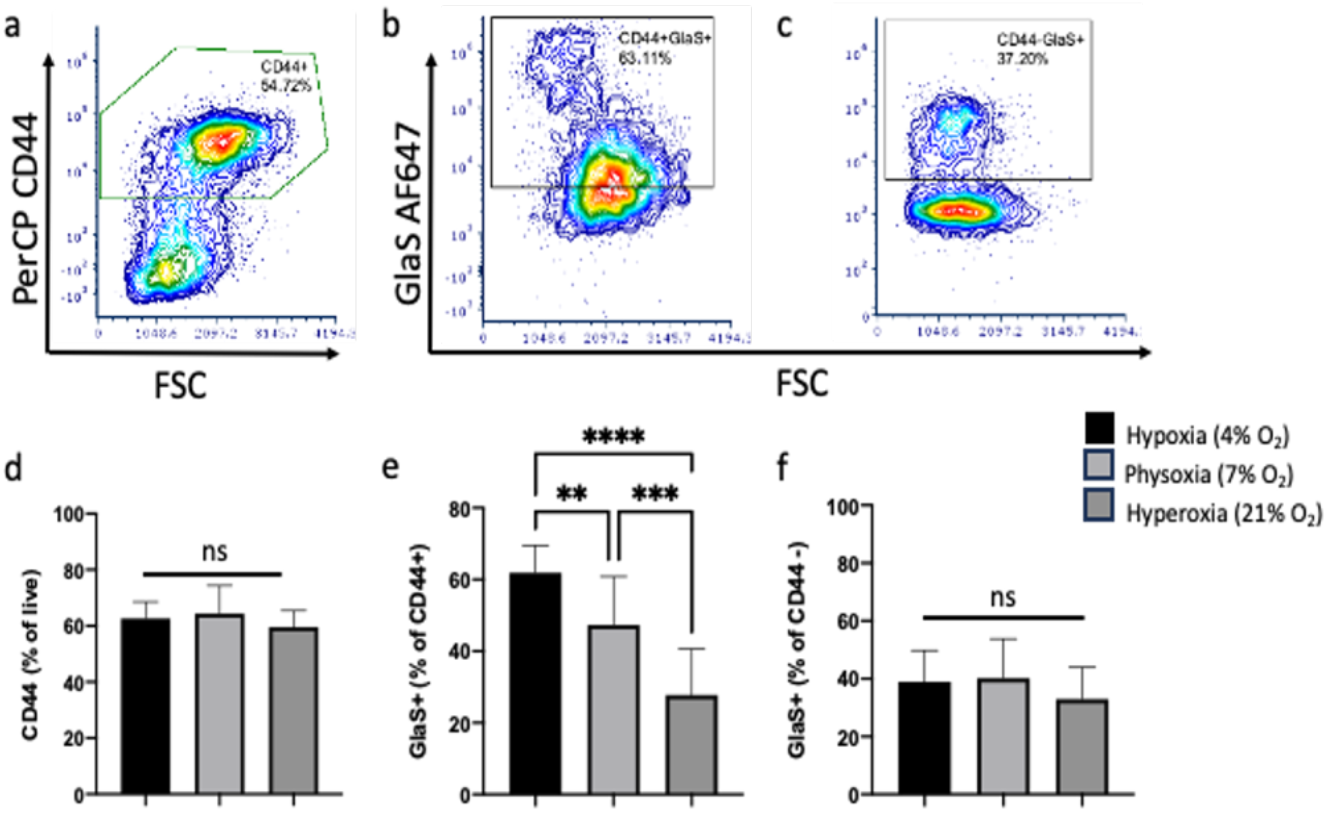
GlaS binding to CD44+ cells is increased in hypoxia. Flow cytometry dot plots of representative (hypoxic) CD44+ (**a**; from live), CD44+GlaS+ (**b**) and CD44-GlaS+ (**c**) populations. No significant difference was found in CD44+ populations between oxygen conditions (**d**). There was significantly more CD44+GlaS+ cells in hypoxic vs physoxic and hyperoxic conditions, and physoxic vs hyperoxic conditions (**e**). There was no significant difference in CD44-GlaS+ populations between oxygen conditions (**f**). n=13 per O^2^ concentration (n=5 PDX tumor types, n=1-3 technical replicates). \*\**p*<.01, \*\*\**p*<.001, \*\*\*\**p*<.0001, ns=not significant

The percent of GlaS+ cells from the live CD44+ population (CD44+GlaS+, **Fig. 1b,e**) were significantly different among all the PDX cell lines at different oxygen conditions (F_2,36_=27.69, *p*<.0001). CD44+GlaS+ frequencies were increased in hypoxic (61%) vs physoxic (47%; *p*=.01) and hyperoxic (28%; *p*<.0001) conditions. The percent of CD44+GlaS+ cells was also significantly increased in the physoxic vs the hyperoxic condition (*p*=.0003). Within a given PDX line, there were significant differences in the frequency of CD44+GlaS+ cells (**Supplementary Fig. 2**) in M1L2T (F_2,6_=67.88, *p*<.0001), M2L2T (*p*=.01) and M3L2G (F_2,6_=37.01, *p*=.0004) cells. There were no differences in the frequency of CD44+GlaS+ cells in CTC092 (F_2,1_=3.85, *p*=.34) and CTC205 (F_2,6_=1.8, *p*=.25) lines. The percentages of GlaS+ cells in the CD44-population (**Fig. 1c,f**) between oxygen conditions were not significantly different (*p*=.054).

**Figure 2.**
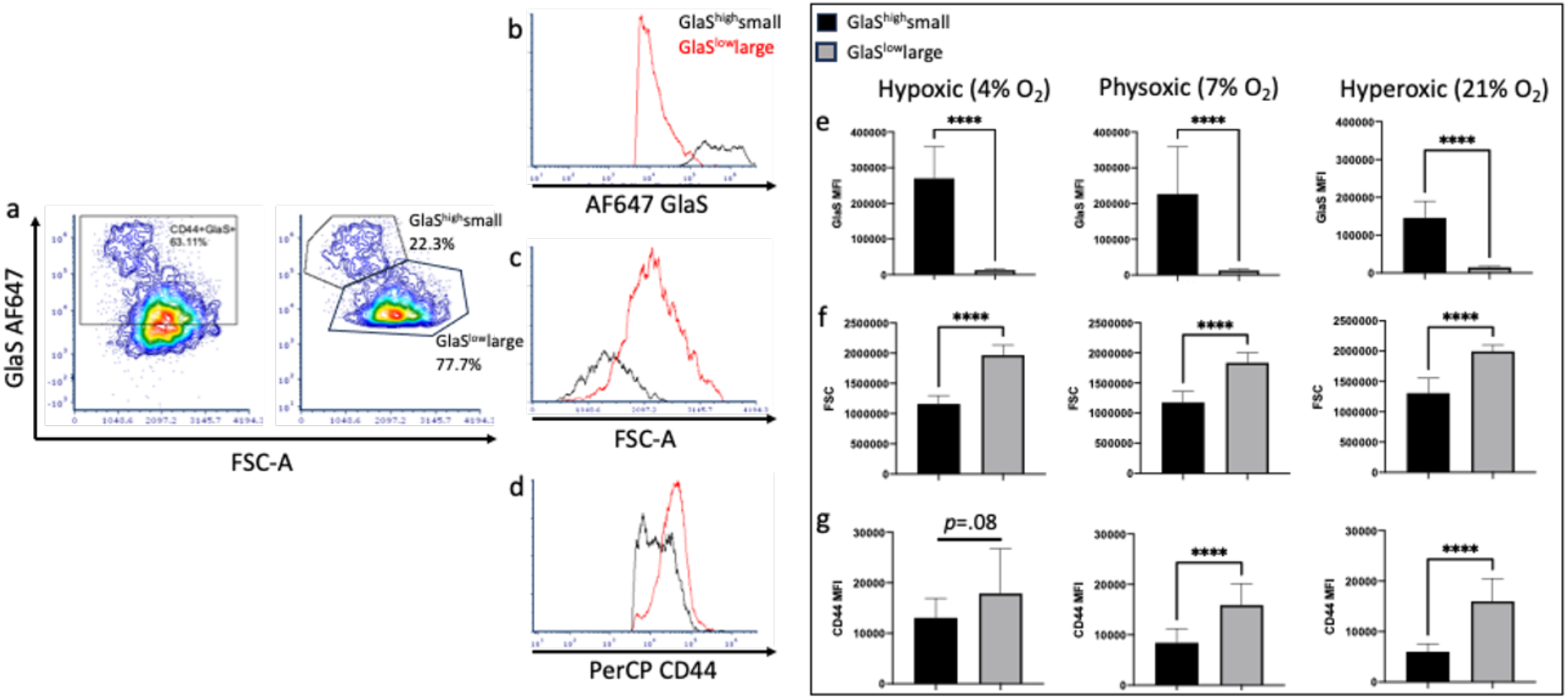
Two populations of CD44+ cells with different sizes, GlaS binding and CD44 expression. Dot plots of flow cytometry data demonstrate that there are CD44+GlaS+, GlaS^high^small and GlaS^low^largepopulations of cells (**a**). Flow cytometry histograms demonstrate AF647 GlaS MFI (**b**), FSC-A (**c**) and PerCP CD44 MFI (**d**) for GlaS^high^ (black) and GlaS^low^ (red) populations. AF647 GlaS MFI (e), FSC-A (f) and PerCP CD44 MFI (g) are quantified for GlaS^high^small and GlaS^low^large populations of cells in hypoxic, physoxic and hyperoxic conditions. n=13 (n=5 PDX tumor types, n=1-3 technical replicates). \*\*\*\**p*<.0001

### The population of smaller-sized CD44+ cells had greater intensity of GlaS staining

From the CD44+ cells, there were two populations which could be distinguished by their size (as determined by forward scatter; FSC) and GlaS staining intensity (as determined by median fluorescence intensity, MFI; **Fig. 2**). The cells with higher FSC (larger) had decreased GlaS MFI (GlaS^low^large) vs a population with lower FSC and increased GlaS MFI (GlaS^high^small). GlaS^low^large cells had lower GlaS binding (**Fig. 2b,e**) and were 1.5x larger (**Fig. 2c,f**) versus the GlaS^high^small cells (*p*<.0001 in all oxygen conditions for FSC (t(24)=13.49; hypoxic, t(24)=9.58; physoxic and t(16.54)=9.72; hyperoxic) and GlaS MFI (t(12.05)=10.45; hypoxic and t(12.11)=10.47; hyperoxic)). GlaS MFI was 21x, 18x and 10x higher in the smaller vs larger CD44+ population at hypoxic, physoxic and hyperoxic conditions, respectively. Differences in GlaS MFI relative to oxygen conditions were apparent in the GlaS^high^small population (*p*=.0008), but not in the GlaS^low^large population (F_2,36_=1.63, *p*=0.21). Further, CD44 MFI was also different in these two populations; cells which were cultured in physoxic (*p*<.0001) and hyperoxic (*p*<.0001) conditions had a higher CD44 MFI in the GlaS^low^large vs the GlaS^high^small population. There were no significant differences in CD44 MFI in the GlaS^low^large vs GlaS^high^small populations in cells cultured under hypoxic conditions (t(16.32)=1.8, *p*=.09).

Although the overall CD44+ population did not change between oxygen conditions, there was a difference in CD44 levels (MFI) when considering the GlaS^low^large and GlaS^high^small populations. Oxygen condition only had an effect on CD44 MFI in the GlaS^high^small population (F_2,36_=19.74, *p*<.0001, **Supplemental Fig. 3a**,**c**), with increased CD44 MFI in the cells cultured in hypoxic conditions vs those cultured in physoxic (*p*=.007) and hyperoxic (*p*<.0001) conditions. Oxygen did not have an effect on the CD44 MFI in the GlaS^low^large cell population (*p*=.96, **Supplemental Fig. 3b**,**d**).

**Figure 3.**
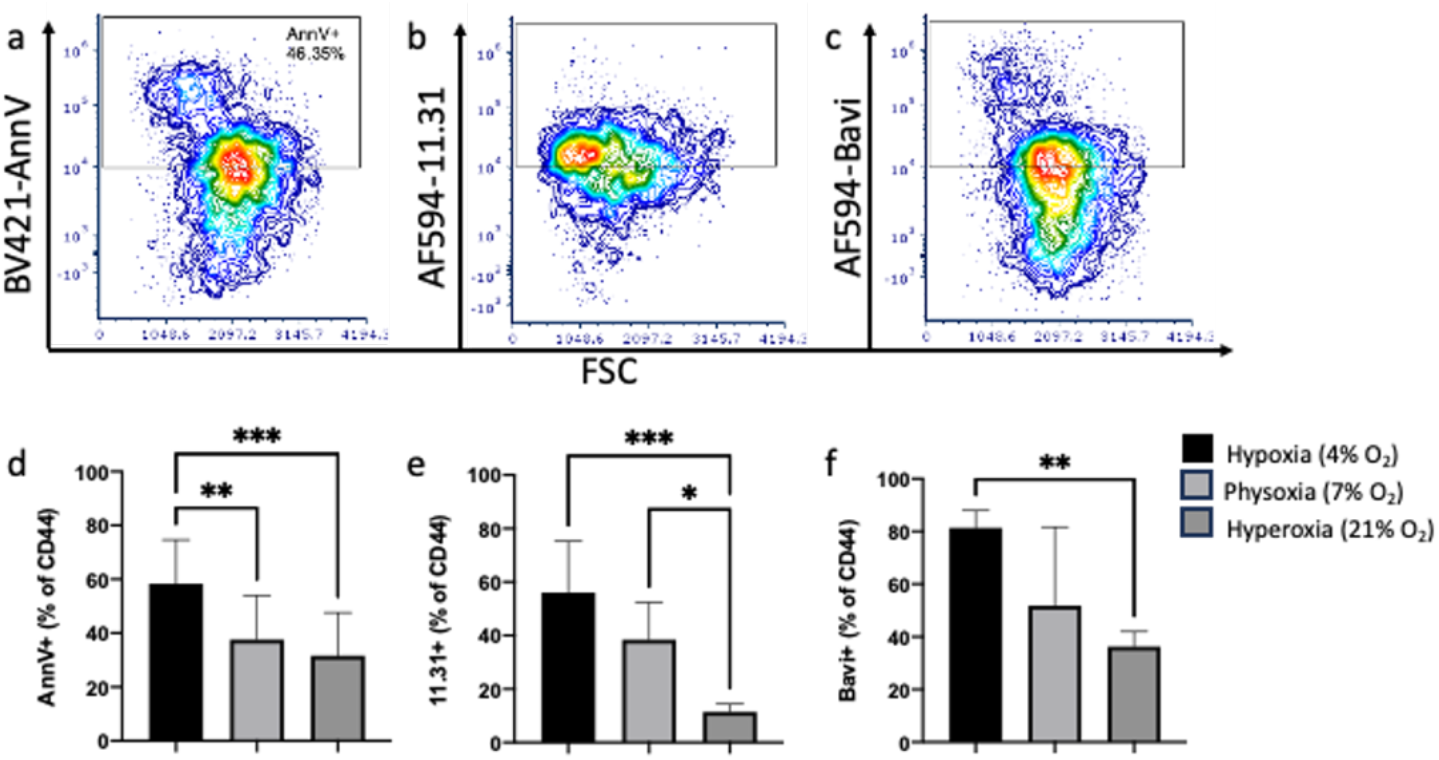
PS-binding by Annexin V, Bavituximab and 11.31 are increased in CD44+ cells in hypoxia. Flow cytometry dot plots of representative (hypoxic) CD44+AnnV+ (**a**), CD44+11.31+ (**b**) and CD44+Bavi+ (**c**) populations. Cells cultured in hypoxic conditions had significantly more AnnV+ (**d**), 11.31+ (**e**) and Bavi+ (**f**) staining vs cells which were cultured in hyperoxic conditions. There was more CD44+AnnV+ cells in hypoxic vs physoxic conditions and more CD44+11.31+ cells in physoxic vs hyperoxic conditions. AnnV; n=13 per O^2^ concentration (n=5 PDX tumor types, n=1-3 technical replicates), Bavi/11.31; n=3-5 per O^2^ concentration (n=3-5 PDX tumor types, n=1 technical replicate). \**p*<.05, \*\**p*<.01, \*\*\**p*<.001

### PS-binding by Annexin V, Antibody 11.31 and Bavituximab on CD44+ cells

PS externalization was confirmed using known PS-binding AnnV as well as antibodies 11.31 and Bavi, validating our findings on GlaS binding. Differences in PS-binding of AnnV, 11.31, and Bavi on CD44+ cells were also observed when cells were held at the different oxygen conditions **(Fig. 3)**. There was a significant difference in AnnV staining among all oxygen conditions (F_2,36_=9.78, *p*=.0004, **Fig. 3d**); specifically, there was significantly more CD44+AnnV+ cells in hypoxic (58%) vs physoxic (37%, *p*=.007) and hypoxic vs hyperoxic (32%, *p*=.0005) conditions. The antibody 11.31 also demonstrated differences in binding between oxygen conditions (*p*=.0009, F_2,12_=13.17, **Fig. 3e**) with an increased frequency of CD44+11.31+ cells in hypoxic (56%) vs hyperoxic (11%, *p*=.0007) and physoxic (39%) vs hyperoxic (*p*=.02) conditions. Bavi binding followed the same trend, with significant differences between all oxygen conditions (F_2,8_=8.3, *p*=.01, **Fig. 3f**) with increased CD44+Bavi+ cells in hypoxic (81%) vs hyperoxic (45%, *p*=.01) conditions.

Although GlaS, AnnV and the PS control antibodies (Bavi and 11.31) all target PS, there were some differences in staining for these four proteins on the PDX lines (**Supplementary Fig. 4**). There were significant differences in PS staining under hypoxic conditions (F_3,31_=3.57, *p*=.025, **Supplementary Fig. 4a**) with more CD44+Bavi+ cells than CD44+AnnV+ (*p*=.023) and CD44+11.31+ (*p*=.035). PS staining was also different among cells cultured under hyperoxic conditions (F_3,31_=3.6, *p*=.024, **Supplementary Fig. 4e)** with the frequency of CD44+11.31+ decreased compared to CD44+AnnV+ (*p*=.029) and CD44+Bavi+ (*p*=.036). There were no significant differences among any of the markers in physoxic conditions (F_3,30_=1.19, *p*=.331, **Supplementary Fig. 4c**). The percentage of CD44+GlaS+ cells were not different from any of the control PS stains under any of the oxygen conditions. However, we observed that GlaS, AnnV and the PS-binding Abs were not always associated with the same cells (**Supplementary Fig. 4**). The percentage of CD44+ cells which were GlaS+, AnnV+ and PS control+ (Bavi or 11.31) were 35%, 22% and 9% for hypoxic, physoxic and hyperoxic (**Supplementary Fig. 4b**,**d**,**f**) conditions, respectively. The remaining cells were GlaS+AnnV+PS control-(9%, 5% and 11%), GlaS+PS control+AnnV-(12%, 12% and 2%), AnnV+PS control+GlaS-(9%, 4% and 2%), GlaS only (6%, 8% and 6%), PS control only (12%, 10% and 7%) and AnnV only (5%, 7% and 9%), for hypoxia, physoxia and hyperoxia, respectively. Overall, the percentages of unstained cells increased as oxygen levels increased (13%, 32% and 54%) for hypoxia, physoxia and hyperoxia, respectively, suggesting that there was a decrease in PS expression as oxygen levels increased resulting in lower PS-associated staining.

**Figure 4.**
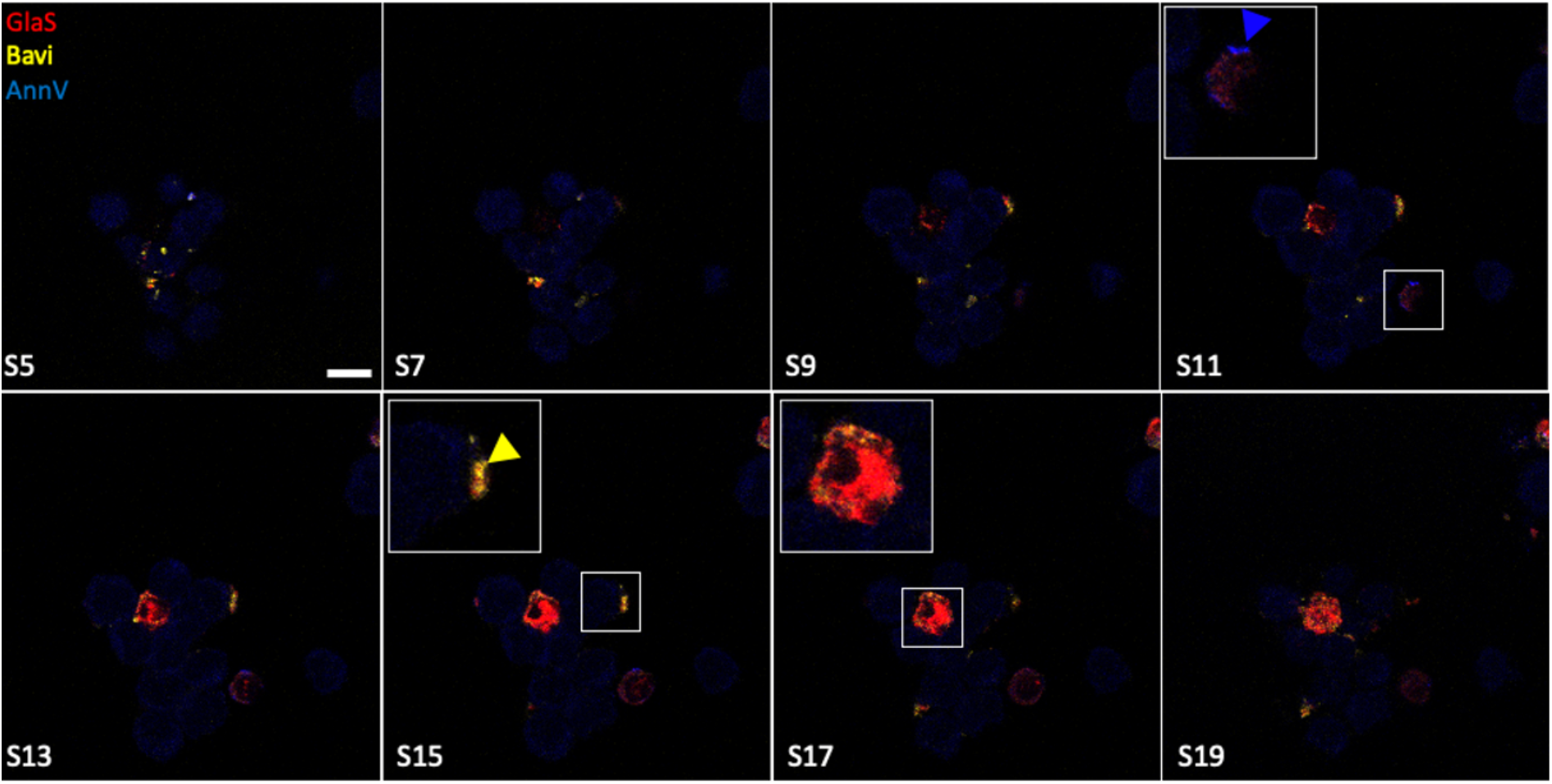
Confocal microscopy of PDX cells reveals intracellular GlaS and cell membrane-bound PS-binding AnnV and Bavi. PDX tumor (M1L2T) was excised from a mouse, dissociated into a single cell suspension and culture overnight at 4% O_2_. Cells were stained with AF647 GlaS (red), AF594 Bavi (yellow) and BV421 Annexin V (blue). Zoomed inset (Section(S)11, S15 and S17). Scale bar = 20 μm.

### Confocal microscopy localizes GlaS to the cytoplasm of PDX derived cells

Microscopy revealed that the localization of GlaS within isolated PDX-derived cells was different from the PS staining controls. When cells are held at hypoxic conditions, GlaS was found both on the membrane of cells as well as within the cells (**Fig. 4, section (S)17 and Supplemental Movie 1**). AnnV (blue arrowhead; **Fig. 4, S11**) and Bavi (yellow arrowhead; **Fig. 4, S15**) were only detected along the membrane of the cells.

### GlaS Accumulates in Hypoxic Regions of Tumors *In Vivo*

Twenty-four hours (h) after intravenous (IV) administration of a combined dose of a hypoxia fluorescent probe, HypoxySense, and HiLight750-GlaS into M1L2T PDX tumor-bearing mice, fluorescence imaging revealed distinct patterns of hypoxia and GlaS accumulation within the tumors (**Fig. 5**). BLI identified viable CSCs throughout the tumor. HypoxySense was used to visualize hypoxic areas, with the tumor cores exhibiting higher levels of hypoxia compared to the surrounding tumor tissue. GlaS accumulated not only within the tumor but also in the gut, knees, ankles, and teeth. In a bisected excised tumor, the localization and spatial resolution of the fluorophores were improved. GlaS localized primarily in the tumor core, where hypoxia was also more pronounced.

**Figure 5.**
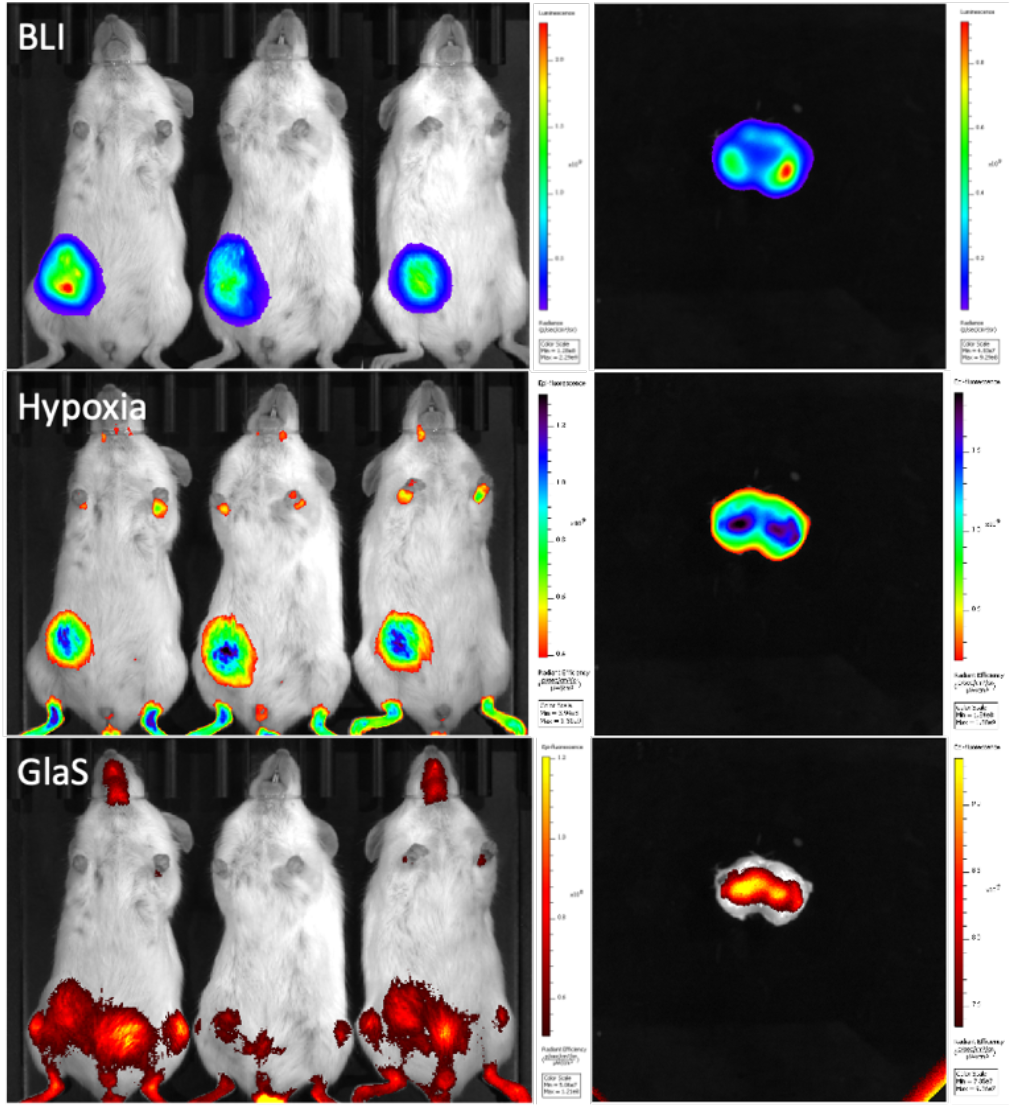
In vivo imaging of hypoxia and GlaS in tumor bearing mice. Bioluminescence imaging (BLI) visualizes viable cancer stem cells (CSC) in M1L2T PDX tumors (n=3). Hypoxia (HypoxySense) and HiLight750-GlaS were visualized using fluorescence imaging, with GlaS found in regions of high hypoxia within the tumor cores.

### GlaS Delivers a Small Molecule to the Cytoplasm of CSC

Bioluminescence was used to monitor the intracellular delivery of luciferin, a small molecule, via GlaS (GlaS-SS-luc). Upon binding to PS, internalization, and reduction of the disulfide bond, luciferin is released and used in the bioluminescent reaction (**Supplementary Fig. 5a**). Differences in the kinetics of light output were observed in culture, when hyperoxic M1L2T cells were incubated with GlaS-SS-luc or equimolar D-luciferin (**Supplementary Fig 5b**). Following incubation with D-luciferin, the peak bioluminescent signal occurred immediately (<2 minutes (mins) post incubation), followed by a decline over ∼20 mins before reaching a signal plateau. In contrast, after incubation with GlaS-SS-luc, bioluminescent signals steadily increased from the initial imaging time point, peaking at 29 mins, with a smaller decrease and subsequent plateau. The area under the curve (AUC) for equimolar free D-luciferin was significantly higher (1.6x, t(3.9)=4, *p*=.018) than that of GlaS-SS-luc (**Supplementary Fig. 5c**), although both signals plateaued similarly. Hypoxic M1L2T cells were not tested in culture, as prolonged exposure to ambient oxygen could have led to inaccurate results.

Similar kinetics trends were observed *in vivo*, with delayed and sustained peak bioluminescent signals following the administration of 50 μg GlaS-SS-luc compared to equimolar free D-luciferin (**Fig 6**). After intraperitoneal (IP) administration of equimolar D-luciferin (**Fig. 6a**), signals peaked at 33 mins, which was similar to the start of the IP GlaS-SS-luc peak (**Fig. 6b**). However, GlaS-SS-luc signals were sustained for 31 mins, while D-luciferin signals declined immediately after peaking. Following intravenous (IV) administration of equimolar D-luciferin (**Fig. 6c**), the peak signal was detected instantly (<1 min post-injection), whereas GlaS-SS-luc showed a delayed peak at 15 mins (**Fig. 6d**). Overall, peak bioluminescent signals were higher after the administration of equimolar D-luciferin compared to GlaS-SS-luc, with 7.5-fold increase for IV and a 30-fold increase for IP administration. AUC was calculated for each treatment, to identify luciferin delivered over that time period (**Fig 6g)**, with significant differences in AUC between the different materials and injection routes (F_3,8_=99.11, *p*<.0001). IV equimolar D-luciferin as well as IV and IP GlaS-SS-luc conjugates delivered significantly lower luciferin (as determined by bioluminescent output) than IP equimolar D-luciferin alone (*p*<.0001 for all). There were no differences when GlaS-SS-luc was administered IP or IV (*p*=.908), nor when GlaS-SS-luc or equimolar D-luciferin were administered IV (*p*=.133).

**Figure 6.**
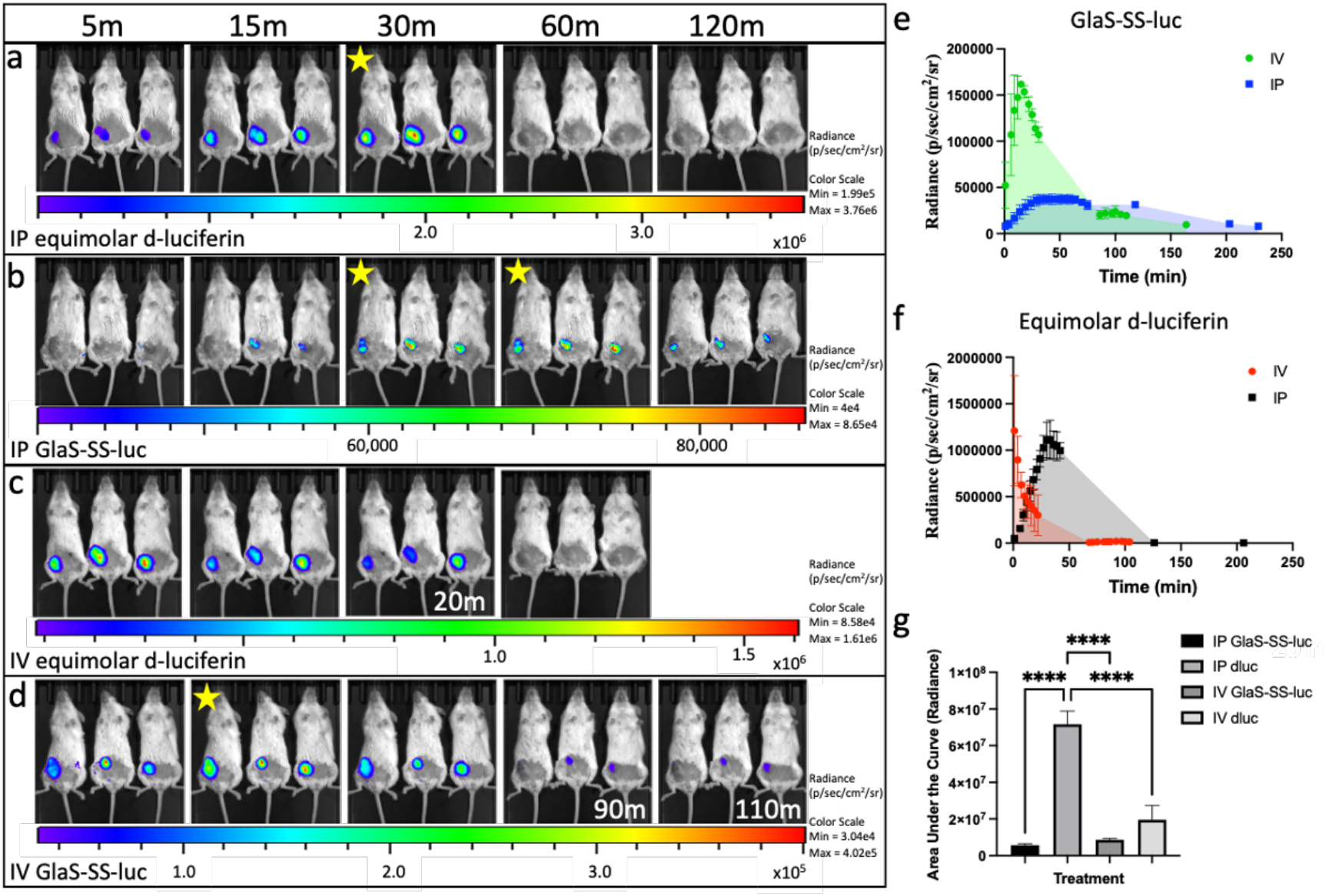
Kinetics of GlaS-SS-luc delivery to cancer stem cells *in vivo*. Mice bearing M1L2T tumors were administered intraperitoneal (IP) equimolar D-luciferin (n=3) or GlaS-SS-luc (n=3, **a&b**) or intravenous (IV) equimolar D-luciferin (n=3) or GlaS-SS-luc (n=3, **c&d**). Mice were imaged over time, with images shown in **a-d** up to 120 minutes (unless noted directly on image). Images with yellow star indicate peak signal after administration of the specific material and injection route. Tumor radiance is demonstrated for GlaS-SS-luc after IP and IV administration (**e**) and equimolar D-luciferin (**f**). Area under the curve is calculated for each treatment (**g**). **** *p*<.0001.

When a lower dose of GlaS-SS-luc (15 μg or ∼0.5 mg/kg) was administered IV into M1L2T PDX tumor-bearing mice, a similar trend was seen at both early (small tumor) and late (large tumor) time points (**Supplemental Fig 6**). In small tumors, signals peaked 7 mins after GlaS-SS-luc administration and immediately (<1 min) after equimolar D-luciferin (**Supplemental Fig 6a**,**b**). There was no significant difference in the AUC between these two materials administered (t(8)=2.26, *p*=.054). Similarly, in larger tumors, signals peaked 7 mins post GlaS-SS-luc and 2 mins after equimolar D-luciferin (**Supplemental Fig 6d**,**e**). In attempt sto increase blood half-life, GlaS was also produced with a Fc fragment (GlaS-Fc) and was conjugated to luciferin (GlaS-Fc-SS-luc) to visualize differences in tumor accumulation and persistence relating to an increase in circulation time (**Supplemental Fig 6f)**. Bioluminescent signals peaked 29 mins post IV administration of GlaS-Fc-SS-luc, showing a delay compared to GlaS-SS-luc. Additionally, AnnV was conjugated to luciferin (AnnV-SS-luc), which appeared to allow for luciferin delivery to the cells, but with no significant peak signal but instead a steady signal (**Supplemental Fig 6g)**. AUC was calculated over a period of 60 mins for each treatment, to identify the luciferin delivered over that time period (**Supplemental Fig 6h)**, with significant differences noted between the four materials administered (*p*<.0001). IV administration of equimolar D-luciferin resulted in the highest AUC as determined by radiance, with GlaS-SS-luc resulting in significantly lower AUC (*p*=.002) whereas the AUC of GlaS-Fc-SSluc and AnnV-SS-luc were not different than D-luciferin (*p*>.99 and *p*=.265, respectively). The AUC of GlaS-SS-luc was not significantly different than those of GlaS-Fc-SS-luc or AnnV-SS-luc (*p*=.265 and *p*>.99, respectively).

## Discussion

The cellular characteristics and location within the hypoxic regions of the TME allow CSCs to evade traditional therapeutics and perpetuate tumor growth even after conventional therapies. As such, novel approaches to eliminating CSC need to be developed that act within the niche that these cells reside. PS exposure has been identified previously on stem cells^21^, as well as healthy vascular endothelial cells, where increased PS externalization was observed after cells were exposed to hypoxic conditions^43^. In the present study, we observed more PS externalization on CSCs under hypoxic conditions and consequently an increase in GlaS binding and internalization into cells. Because CSCs reside within low oxygen environments *in vivo*, it was anticipated that there would be increased PS externalization when compared to other regions of the tumor, or to cells from normal, healthy tissue. These observations suggest that GlaS may provide an opportunity to target the enhanced PS on the outside of the CSCs for directed delivery to cytoplasmic and nuclear targets within these cells. This would enable delivery of therapies (e.g. small molecules, radiation and/or biologics) directed at intracellular pathways with limited off-target effects. Further, if the externalization of PS is universally found on all CSC, GlaS could be used to treat CSC in any tumor type.

AnnV and Bavi are well known proteins used to bind PS. AnnV is commonly used to identify cells undergoing apoptosis, by specifically targeting PS due to loss of plasma membrane asymmetry^44^. Bavi is a mAb which acts as an immune modulating therapeutic, used to target PS exposed on the surface of tumor cells in the presence of β 2 glycoprotein (β2GPI)^45^. The antibody 11.31 (PGN632) is another PS-targeting mAb which does not require the presence of β2GPI, and has direct binding to PS^46,47^. These three characterized PS-binding proteins were used to confirm our observations of GlaS binding to CD44+ breast CSC. GlaS was found on the same percentage of CD44+ cells as AnnV, Bavi and 11.31, validating that it binds externalized PS. However, these PS binding proteins did not always appear bound to the same cells, as determined by flow cytometry and microscopy^21^ suggesting there may be differences in binding requirements for each of these PS-binding proteins. The cellular localization of these labeled proteins was examined using confocal microscopy, and only GlaS appeared to be internalized into the PDX cells. AnnV and Bavi appeared bound only to the cell membrane. AnnV has been observed previously to be internalized by apoptotic cells as well as in HeLa immortalized human cells, identified within vesicles^48^. However, in live cells, this has not been observed in our studies^21^. Since the fluorophores used in these studies are small molecules, similar in size to chemotherapeutic agents, our observations reveal an opportunity to deliver molecular payloads into CSC and target intracellular pathways such as those related to Ras and other oncogenes (reviewed in^49^). In contrast, Bavi bound to exposed PS but remained on the outer surface of cells which may be useful for immune modulation^29^, but would not enable targeting of intracellular pathways.

The kinetics of GlaS-mediated internalization and its ability to functionally deliver a small molecule into the cytoplasm were studied using luciferin as a drug mimetic. Although the presence of O_2_ is required in the bioluminescent reaction, even in extreme hypoxia (∼0% 0_2_) light can be produced^50^. These changes in O2 concentration do influence signal intensities, with lower O2 producing lower light output^50^. Since the CD44+ breast CSCs were engineered to express luciferase, and the luciferase reaction requires intracellular luciferin delivery, this system acts as a molecular light switch to track the localization of GlaS-conjugates and confirm the functional delivery of molecular cargo to CSCs both in culture and *in vivo*. ^50^In both settings, bioluminescent signals—indicating luciferin delivery—were delayed and sustained when GlaS-SS-luc was administered, compared to equimolar free D-luciferin. D-luciferin can diffuse readily through the cell membrane, leading to a rapid bioluminescent response, whereas GlaS-SS-luc must first bind to PS and be internalized, followed by disulfide bond reduction to release luciferin, enabling the bioluminescent reaction. These distinct mechanisms likely contribute to the observed differences in bioluminescence kinetics. In attempts to increase blood half-life of GlaS, a GlaS was produced with an Fc region, known to increase the serum half-life of therapeutic antibodies^51–53^. The addition of the Fc region changed the dynamics of luciferin delivery, as determined by bioluminescence, when compared to GlaS alone. However, there was no significant difference in the overall delivery of the luciferin over time. Given that there was no significant benefit, the GlaS protein is adequate in cargo delivery to the CSC. The use of AnnV-SS-luc was intended to be a control, with our previous studies indicating that when attached to a fluorophore, it remains on the outside of the cell membrane. Interestingly, when AnnV-SS-luc was administered, we did observe bioluminescence, indicating the intracellular presence of the luciferin cargo. Here, the luciferin was either internalized while still conjugated to AnnV, followed by release of the luciferin, or the SS bond was broken outside of the cell followed by passive diffusion of the luciferin into the cell alone, either resulting in bioluminescence signals.

CD44 expression has been identified in many solid malignancies and is a well-known marker of CSCs^54–57^. In breast cancer, CD44+ CSCs are shown to exhibit high tumorigenicity and the ability to establish tumor heterogeneity following transplantation into mice^57^. An increase in CD44 expression identified in cells cultured under hypoxia has been observed before in breast cancer cell lines^58–60^ as well as other models of cancer including glioma^61,62^, gastric^10^ and ovarian^63^. In this study, we observed that CD44 expression in the GlaS^high^small cells was sensitive to oxygen condition in which the cells were cultured. Although a marker of CSC after lineage depletion, CD44 is expressed on normal cells including lymphocytes, fibroblasts, and smooth muscle cells in both humans and mice^64^ making it a complicated target for directed delivery to the CSC.

Typically, CSCs reside within hypoxic areas of the tumor and can be identified based on their size; small sized cells are considered to be the quiescent, slow cycling stem cells^65–67^. Further, smaller sized CSCs have been found to enhance tumorigenicity *in vivo*^68^. Our study found that GlaS better targeted the small CD44+ cells (GlaS^high^small), and further, had enhanced targeting under hypoxic conditions. These findings have significant implications, suggesting that GlaS could serve as a therapeutic delivery tool to specifically target the challenging-to-treat CSCs residing in the hypoxic TME.

Difficulties in targeting and treating CSCs have led to tumor relapse and poor patient outcomes. In this study, we reveal the natural PS externalization on the CSCs which is enhanced under hypoxia and demonstrate intracellular delivery following GlaS binding to PS. GlaS targeted CD44+ cells, with superior targeting of the small CD44+ population, which was likely the small, tumorigenic CSC residing in the hypoxic TME. In the future, GlaS can serve as a platform for a PS-mediated intracellular targeted therapeutic delivery to potentially treat all tumor CSCs in their native hypoxic regions.

## Methods

### Patient derived xenografts

Human patient derived xenografts (PDX)s^40–42^ were derived by, and labeled by, Dr. Huiping Liu and her team (Northwestern University). Cells were derived from human triple negative breast cancer. Previously, CD44+ lineage negative CSC were isolated and engineered to express luciferase2- and tdtomato (L2T) or eGFP (L2G) using lentivirus transduction^40^. For flow cytometry to determine GlaS staining under variable oxygen conditions, five PDX lines were used, which included M1L2T, M2L2T, M3L2G, CTC205 (L2T) and CTC092 (L2T). For GlaS-mediated luciferase delivery, the M1L2T PDX line was used due to ease of growth. Cells were expanded in NOD-scid gamma (NSG) mice, obtained from Jackson laboratories (age 6-8 weeks). Cells from frozen samples were resuspended in PBS, mixed 1:1 with Matrigel to a total volume of 50 μl, and were injected into the right lower mammary fat pad. Because tumors varied in growth rate (ready to harvest in 2-6 months), when all PDX lines were to be used at the same time, injections were staggered so that the tumors were approximately the same size when excised. When cells were used in culture, they were maintained in HuMEC media (Fisher Sci, cat#12-752-010) with growth factors (bovine pituitary extract and HuMEC supplements, added as per manufacturer’s instructions), 5% FBS and 0.5% penicillin/streptomycin. Cells were incubated at 37°C, 5% CO_2_ and 4% O_2_ (hypoxic), 7% O_2_ (physoxic) or 21% O_2_ (hyperoxic). To collect cells, supernatant was set aside, plates were incubated with trypsin for 5 min at 37°C followed by gentle scraping. Plates were washed using PBS and added to supernatant to collect cell pellet by centrifugation.

### Expression and Purification of GlaS and GlaS-Fc protein and GlaS fluorescent conjugation

GlaS protein was expressed and purified as described^21^. For fluorescence conjugation, purified protein underwent centrifugal ultrafiltration, TCEP reduction and maleimide conjugation to AlexaFluor (AF)647. AF647 GlaS was analyzed using size-exclusion chromatography and UV-Vis.

GlaS-Fc is a genetic fusion protein comprised of the amino terminal of human Protein S (Gla, TSR, EGF1-4 domains) linked to the murine IgG2A CH2 and CH3 domains via an eight amino acid linker domain (EPRGPTIK) derived from the IgG hinge region (**Supplemental Figure 7**). The Fc domain contains a single amino acid substitution (IgG2A I253A) to minimize binding to FCRn. Exact molecular weight of GlaS-Fc is 53000.7461 Da and pI is estimated to be 5.30. GlaS-Fc protein was stably expressed in CHO K1 cells and grown in roller bottles seeded with a stable cell pool at 3 × 10^6^ cells/ml in media supplemented with 60 ng/ml vitamin K required for post-translational gamma-carboxylation of glutamic acids within the N-terminal Gla domain. Media was harvested, clarified by centrifugation, concentrated 10X and diafiltered into Tris Buffered Saline (TBS) containing 300mM Betaine and frozen until purification could be initiated. Production of Gla-Fc was monitored by anti Gla Western blot. Purification of the GlaS-Fc protein entailed a three-step process. Thawed media was first loaded onto a Protein G column which binds the Fc domain. The column was then washed with TBS containing 0.5M NaCl to remove weakly bound proteins and then eluted with pH 5 glycine. Eluted fractions were immediately neutralized with Tris. GlaS-Fc containing fractions were pooled, diluted, adjusted to pH 5.5 and loaded onto a Sulfate column. After washing to baseline in 50mM NaCL pH 5.5, the column was eluted with a linear gradient of 50mM - 1 M NaCl, pH 5.5. GlaS-Fc containing fractions were pooled and characterized for purity by SE-HPLC and SDS-PAGE. Following concentration to 3 ml, preparative SEC on SuperDex 200 was run final polishing step. Mobile phase was 20mM HEPES, 150mM NaCl, 10mM CaCl_2_, pH 7.2. The final pooled fractions were pooled, 1% sucrose added and stored frozen at -80°C.

### Conjugation of GlaS and GlaS-Fc to luciferin

A solution of the protein in a storage buffer (1 mg/ml in HEPES 20 mM, NaCl 150 mM, CaCl_2_ 10 mM pH 7.2) was treated with TCEP (4 equiv., 10 mM in DI water) for 4 h at +4°C. A TTL thiol labeling luciferin reagent (VitaLume Biotechnologies, Cat# VB10302; 10 equiv., 10 mM in DMSO) was added to the solution and the resulting mixture was incubated for 20 h at +4°C. The product was filtered into the same storage buffer through a spin desalting column (0.5 ml, Zeba ™, 7k MWCO). GlaS-SS-Luc and GlaS-Fc-SS-Luc conjugates were characterized using UV-Vis measurements and SEC.

### Conjugation of AnnV to luciferin

A solution of AnnV in water (1 mg/ml) was treated with an ATL amine labeling luciferin reagent (VitaLume Biotechnologies. Cat# VB10301; 10 equiv., 10 mM in DMSO) and mixed. After incubation for 20 h at +4°C, the product was filtered into the storage buffer through a spin desalting column (0.5 ml, Zeba ™, 7k MWCO). AnnV-SS-luc was characterized using UV-Vis measurements and SEC.

### Tumor digestion

Tumors were excised, minced into ∼2 mm pieces and incubated in a digestion solution (100 U/ml DNAse I (Stem Cell, cat#7900), Collagenase/Hyaluronidase 1X (Stem Cell, cat#7912) in RPMI) for 60 min, on a shaker at 80 rpm in a humidified incubator at 37°C and 5% CO_2_. Cells were then filtered through a 70 μm strainer using PBS to wash plate and filter. Cells were centrifuged for 5 mins at 300G, supernatant aspirated and resuspended in 2 ml ACK lysis buffer for 1 min. 9 ml PBS + 10% FBS was then added and suspension was centrifuged again. Cells were resuspended in PBS followed by a sample being used for cell counting and viability using the trypan blue assay (Countess automated cell counter; Invitrogen, CA USA).

### Flow cytometry

Following tumor digestion (above), cells were seeded into 75 mm flasks (n=3 for each PDX line) at a confluency of with 3×10^6^-10×10^6^ cells total, depending on total number collected from tumors. One flask from each cell line was incubated at 4%, 7% or 21% O_2_ at 37°C and 5% CO_2_ overnight (∼20 hours). After overnight incubation in prescribed oxygen levels, cells were collected by supernatant, trypsinization, scraping and washing with PBS. All collection and staining procedures were performed in the appropriate oxygen condition. Cells were centrifuged at 300G for 5 mins when they were counted for staining. 1×10^6^ cells were seeded into a 96-well round bottom plate. All staining steps were performed in 100 μL volume in the dark. Samples were first incubated with Zombie NIR viability dye (1:750, BioLegend) for 30 min. Cells were washed once with flow buffer, followed by incubation with TruStain FcX PLUS (anti-mouse CD16/32) Antibody (BioLegend, Cat#156603; 0.25 μg/sample) and Human TruStain FcX (BioLegend, Cat#422301; 5 μl/sample) for 10 min. AF647 GlaS (1 μg/sample), PerCP CD44 (BioLegend, Cat#103035; 0.125 μg/sample) and AF594 Bavi or 11.31 antibodies (Birge Lab, Rutgers), each at 1 μg/sample, were then added and incubated for 30 min at room temperature. Bavi and 11.31 were used as PS-binding controls and stained with Alexa Fluor™ 594 Microscale Protein Labeling Kit (ThermoFisher, cat#A30008). Cells were washed twice with flow buffer prior to staining with BV421 Annexin V (BioLegend, Cat#141703; 2.5 μl/sample) in Annexin binding buffer (BioLegend, Cat#422201). Cells were washed twice with flow staining buffer and fixed with 4% PFA for 10 min and resuspended in a final volume of 100 μL for flow cytometry analysis using the Cytek Aurora spectral flow cytometer (Cytek, CA, USA). Single stained controls and unstained controls for all conditions were used to assess fluorescent spread and for gating strategies. Flow cytometry data was analyzed with the software FCSExpress (DeNovo Software, CA, USA). The data presented herein were obtained using instrumentation in the MSU Flow Cytometry Core Facility. The facility is funded in part through the financial support of Michigan State University’s Office of Research & Innovation, College of Osteopathic Medicine, and College of Human Medicine.

### In Vivo Fluorescence Imaging of GlaS and Hypoxia Localization in PDX tumors

M1L2T tumors were grown in NSG mice for 32 days. M1L2T was used as this is the most consistent and fastest growing PDX cell line. GlaS was conjugated to HiLight Fluor 750 (AnaTag™ HiLyte™ Fluor 750 Microscale Protein Labeling Kit, cat# AS-72044) as per manufacturer’s protocol. A combined 2 nmol IVISense Hypoxia CA IX 680 Fluorescent Probe (HypoxySense, Revvity, cat#NEV11070) and 30 μg HiLight Fluor 750-GlaS were injected iv into mice. 24 h post injection, mice were injected ip with 100 μl (30 mg/ml) d-luciferin, anesthetized, and imaged using the Spectrum In Vivo Imaging System (IVIS, PerkinElmer) following fluorescence filter sets: HypoxySense (Ex 675/Em 720) and HiLight Fluor 750-Glas (Ex 745/Em 800). BLI was performed approximately 5 min post d-luciferin injection. One tumor was excised, cut in half and imaging was performed as above. Tumors were fixed in 4% PFA for microscopy.

### GlaS-SS-luc Delivery in Culture

M1L2T tumors were grown in NSG mice, excised, and dissociated into a single cell suspension (as above). Cells were plated at 5×10^4^ cells/well in a 96 well plate and maintained in hyperoxic conditions for 24h prior to the addition of 1 μg GlaS-SS-luc or equimolar free d-luciferin (PerkinElmer, cat#122799). Hypoxic conditions were not tested due to exposure to higher oxygen levels when taken from the hypoxic incubator for imaging. Immediately following addition, bioluminescence (radiance: photons/sec/cm^2^/sr) from the cells was imaged over a period of 2 h, using the IVIS Lumina.

### GlaS-SS-luc, GlaS-Fc-SS-luc and AnnV-SS-luc Delivery to Tumors In Vivo

NSG mice bearing M1L2T tumors were used to validate intracellular delivery of the small molecule, luciferin, *in vivo*. First, route of administration (intravenous; IV or intraperitoneal; IP) was tested. Mice were anesthetized using 1.5-2% isoflurane in O2 followed by the administration of 50 μg of GlaS-SS-luc conjugate (n=3 IP and n=3 IV) or equimolar d-luciferin (n=3 IP and n=3 IV) Mice were immediately placed into the IVIS for bioluminescent imaging (BLI) for up to 200 min. To examine any differences in luciferin delivery in different sized tumors, mice were anesthetized using 1.5-2% isoflurane in O2 followed by the IV administration of 15 μg GlaS-SS-luc conjugate (n=5, small tumors and n=5, large tumors), equimolar d-luciferin (n=5, small tumors and n=5, large tumors), GlaS-Fc-SS-luc (n=3, large tumors) and AnnV-SS-luc (n=3, large tumors). GlaS-Fc-SS-luc and AnnV-SS-luc were injected at a dose which contained equimolar luciferin molecules in GlaS-SS-luc. Mice were immediately placed into the IVIS for BLI for up to 70 min.

### Stastical Analysis

Statistics were performed using PRISM software (GraphPad, MA, USA; software version 9.0.0). Data is presented as mean +/-standard deviation (sdev). Staining from hypoxic, physoxic and hyperoxic conditions and AUC values from bioluminescence assays were compared using an unpaired t-test (comparing 2 conditions) or a one-way ANOVA with Tukey’s multiple comparisons test (comparing 3 or more conditions). Equal variances were tested using Brown-Forsythe test and normal distribution was tested using Shapiro-Wilk test. If data failed the equal variances test, a Welch’s ANOVA test with Dunnet’s T3 multiple comparisons test was performed. If data failed the normality test, the non parametric Kruskal-Wallis with Dunn’s multiple comparisons test was performed. Staining from GlaS^high^small and GlaS^low^large cells were compared using a two-tailed unpaired t test. If the data failed the D’Agostino & Pearson normality test, a nonparametric two-tailed Mann-Whitney test was performed. If the data did not have equal variances (F test), an unpaired two-tailed t test with Welch’s correction was performed. A finding of *p*<.05 was considered significant.

### Microscopy

M1L2T cells dissociated from a tumor were placed into hypoxic conditions overnight. Cells were then incubated with AF647 GlaS, BV421 AnnV and AF594 Bavi for 30 mins at room temperature. Cells were washed twice with staining buffer prior to fixation using 4% PFA for 10 min. A sample of 200,000 cells were spun onto a slide using a cytospin followed by placement of a coverslip for imaging. Imaging was performed using the Leica TCS SP8 X spectral upright confocal microscope system. Voxel size (x,y,z): 0.17 × 0.17 × 0.5 μm^3^. Images were prepared using Fiji software (ImageJ; V2.9.0)^69^.

## Author contributions

AVM, TT, TH and CHC conceptualized the study. AVM and TT performed experiments, data analysis and statistics. AVM, TT, TH and CHC contributed to the analysis and/or interpretation of the data. DS and TH helped design the GlaS and GlaS-Fc. HL provided the PDX cell lines and technical advice surrounding growth and maintenance. PK and EG developed the GlaS-SS-luc, GlaS-Fc-SS-luc and AnnV-SS-luc materials and provided guidance on their use. AVM drafted the manuscript. All authors revised and edited the manuscript.

## Acknowledgements

We thank the MSU Flow Cytometry Core for their assistance in experimental planning, acquisition and analysis. We also thank the IQ Microscopy Core for their assistance in confocal microscopy training and help with acquisition and IQ Imaging Core for their support in imaging studies. This work was supported in part by GLAdiator Biosciences and The James and Kathleen Cornelius Endowment (to CHC). This work is partially supported by NIH/NCI R01CA245699 and American Cancer Society ACS0137006 (H.L.).

## References

1. Shiozawa, Y., Nie, B., Pienta, K. J., Morgan, T. M. & Taichman, R. S. Cancer stem cells and their role in metastasis. Pharmacol Ther 138, 285–293 (2013).

2. Singh, S. K. et al. Identification of human brain tumour initiating cells. Nature 432, 396–401 (2004).

3. Al-Hajj, M., Wicha, M. S., Benito-Hernandez, A., Morrison, S. J. & Clarke, M. F. Prospective identification of tumorigenic breast cancer cells. Proceedings of the National Academy of Sciences 100, 3983–3988 (2003).

4. Lapidot, T. et al. A cell initiating human acute myeloid leukaemia after transplantation into SCID mice. Nature 367, 645–648 (1994).

5. Soltanian, S. & Matin, M. M. Cancer stem cells and cancer therapy. Tumor Biology 32, 425–440 (2011).

6. Yang, L. et al. Targeting cancer stem cell pathways for cancer therapy. Signal Transduct Target Ther 5, 8 (2020).

7. McKeown, S. R. Defining normoxia, physoxia and hypoxia in tumours—implications for treatment response. Br J Radiol 87, 20130676 (2014).

8. Mas-Bargues, C. et al. Relevance of Oxygen Concentration in Stem Cell Culture for Regenerative Medicine. Int J Mol Sci 20, (2019).

9. Heddleston, J. M., Li, Z., McLendon, R. E., Hjelmeland, A. B. & Rich, J. N. The hypoxic microenvironment maintains glioblastoma stem cells and promotes reprogramming towards a cancer stem cell phenotype. Cell cycle 8, 3274–3284 (2009).

10. Liang, G. et al. Hypoxia regulates CD44 expression via hypoxia-inducible factor-1α in human gastric cancer cells. Oncol Lett 13, 967–972 (2017).

11. Holmquist-Mengelbier, L. et al. Recruitment of HIF-1a; and HIF-2a; to common target genes is differentially regulated in neuroblastoma: HIF-2a; promotes an aggressive phenotype. Cancer Cell 10, 413–423 (2006).

12. Bhaskara, V. K., Mohanam, I., Rao, J. S. & Mohanam, S. Intermittent hypoxia regulates stem-like characteristics and differentiation of neuroblastoma cells. PLoS One 7, e30905 (2012).

13. Lendeckel, U. & Wolke, C. Redox-Regulation in Cancer Stem Cells. Biomedicines 10, (2022).

14. Heddleston, J. M., Li, Z., McLendon, R. E., Hjelmeland, A. B. & Rich, J. N. The hypoxic microenvironment maintains glioblastoma stem cells and promotes reprogramming towards a cancer stem cell phenotype. Cell cycle 8, 3274–3284 (2009).

15. Nascimento-Filho, C. H. V et al. From Tissue Physoxia to Cancer Hypoxia, Cost-Effective Methods to Study Tissue-Specific O2 Levels in Cellular Biology. Int J Mol Sci 23, 5633 (2022).

16. Hankins, H. M., Baldridge, R. D., Xu, P. & Graham, T. R. Role of flippases, scramblases and transfer proteins in phosphatidylserine subcellular distribution. Traffic 16, 35–47 (2015).

17. Fadok, V. A. et al. Exposure of phosphatidylserine on the surface of apoptotic lymphocytes triggers specific recognition and removal by macrophages. J Immunol 148, 2207–2216 (1992).

18. Srivasatava, K. R., Majumder, R., Kane, W. H., Quinn-Allen, M. A. & Lentz, B. R. Phosphatidylserine and FVa regulate FXa structure. Biochemical Journal 459, 229–239 (2014).

19. Riedl, S. et al. In search of a novel target—Phosphatidylserine exposed by non-apoptotic tumor cells and metastases of malignancies with poor treatment efficacy. Biochimica et Biophysica Acta (BBA)-Biomembranes 1808, 2638–2645 (2011).

20. Vallabhapurapu, S. D. et al. Variation in human cancer cell external phosphatidylserine is regulated by flippase activity and intracellular calcium. Oncotarget 6, 34375 (2015).

21. Hardy, J. et al. Gla-domain mediated targeting of externalized phosphatidylserine for intracellular delivery. The FASEB Journal 37, e23113 (2023).

22. Jaiswal, S. et al. CD47 is upregulated on circulating hematopoietic stem cells and leukemia cells to avoid phagocytosis. Cell 138, 271–285 (2009).

23. Riedl, S. et al. In search of a novel target—Phosphatidylserine exposed by non-apoptotic tumor cells and metastases of malignancies with poor treatment efficacy. Biochimica et Biophysica Acta (BBA)-Biomembranes 1808, 2638–2645 (2011).

24. Birge, R. B. et al. Phosphatidylserine is a global immunosuppressive signal in efferocytosis, infectious disease, and cancer. Cell Death Differ 23, 962–978 (2016).

25. Wang, W. et al. Mobilizing phospholipids on tumor plasma membrane implicates phosphatidylserine externalization blockade for cancer immunotherapy. Cell Rep 41, (2022).

26. Van Rite, B. D. & Harrison, R. G. Annexin V-targeted enzyme prodrug therapy using cytosine deaminase in combination with 5-fluorocytosine. Cancer Lett 307, 53–61 (2011).

27. Guillen, K. P., Ruben, E. A., Virani, N. & Harrison, R. G. Annexin-directed β-glucuronidase for the targeted treatment of solid tumors. Protein Engineering, Design and Selection 30, 85–94 (2017).

28. Gerber, D. E. et al. Tumor-Specific Targeting by Bavituximab, a Phosphatidylserine-Targeting Monoclonal Antibody with Vascular Targeting and Immune Modulating Properties, in Lung Cancer Xenografts. Am J Nucl Med Mol Imaging vol. 5 www.ajnmmi.us/ (2015).

29. Yin, Y., Huang, X., Lynn, K. D. & Thorpe, P. E. Phosphatidylserine-targeting antibody induces M1 macrophage polarization and promotes myeloid-derived suppressor cell differentiation. Cancer Immunol Res 1, 256–268 (2013).

30. Hardy, J. et al. Gla-domain mediated targeting of externalized phosphatidylserine for intracellular delivery. bioRxiv 2022.06.13.495901 (2022) doi:10.1101/2022.06.13.495901.

31. Onzi, G., Guterres, S. S., Pohlmann, A. R. & Frank, L. A. Passive Targeting and the Enhanced Permeability and Retention (EPR) Effect. in The ADME Encyclopedia (2021). doi:10.1007/978-3-030-51519-5_108-1.

32. Shinde, V. R., Revi, N., Murugappan, S., Singh, S. P. & Rengan, A. K. Enhanced permeability and retention effect: A key facilitator for solid tumor targeting by nanoparticles. Photodiagnosis and Photodynamic Therapy vol. 39 Preprint at 10.1016/j.pdpdt.2022.102915 (2022).

33. Tiwari, H. et al. Recent Advances in Nanomaterials-Based Targeted Drug Delivery for Preclinical Cancer Diagnosis and Therapeutics. Bioengineering vol. 10 Preprint at 10.3390/bioengineering10070760 (2023).

34. Attia, M. F., Anton, N., Wallyn, J., Omran, Z. & Vandamme, T. F. An overview of active and passive targeting strategies to improve the nanocarriers efficiency to tumour sites. Journal of Pharmacy and Pharmacology vol. 71 Preprint at 10.1111/jphp.13098 (2019).

35. Fu, Z., Li, S., Han, S., Shi, C. & Zhang, Y. Antibody drug conjugate: the “biological missile” for targeted cancer therapy. Signal Transduction and Targeted Therapy vol. 7 Preprint at 10.1038/s41392-022-00947-7 (2022).

36. Gonda, K. et al. Heterogeneous Drug Efficacy of an Antibody-Drug Conjugate Visualized Using Simultaneous Imaging of Its Delivery and Intracellular Damage in Living Tumor Tissues. Transl Oncol 13, (2020).

37. Van de Wiele, C. et al. Apoptosis imaging in oncology by means of positron emission tomography: A review. International Journal of Molecular Sciences vol. 22 Preprint at 10.3390/ijms22052753 (2021).

38. Mandl, S. J. et al. Multi-modality imaging identifies key times for annexin V imaging as an early predictor of therapeutic outcome. Mol Imaging 3, (2004).

39. Blankenberg, F. G. In vivo imaging of apoptosis. Cancer Biology and Therapy vol. 7 Preprint at 10.4161/cbt.7.10.6934 (2008).

40. Liu, H. et al. Cancer stem cells from human breast tumors are involved in spontaneous metastases in orthotopic mouse models. Proceedings of the National Academy of Sciences 107, 18115–18120 (2010).

41. Tsai, C.-F. et al. Surfactant-assisted one-pot sample preparation for label-free single-cell proteomics. Commun Biol 4, 265 (2021).

42. Dashzeveg, N. K. et al. Dynamic glycoprotein hyposialylation promotes chemotherapy evasion and metastatic seeding of quiescent circulating tumor cell clusters in breast cancer. Cancer Discov CD–22 (2023).

43. Ran, S. & Thorpe, P. E. Phosphatidylserine is a marker of tumor vasculature and a potential target for cancer imaging and therapy. International Journal of Radiation Oncology*Biology*Physics 54, 1479–1484 (2002).

44. Vermes, I., Haanen, C., Steffens-Nakken, H. & Reutellingsperger, C. A novel assay for apoptosis flow cytometric detection of phosphatidylserine expression on early apoptotic cells using fluorescein labelled annexin V. J Immunol Methods 184, 39–51 (1995).

45. Soares, M. M., King, S. W. & Thorpe, P. E. Targeting inside-out phosphatidylserine as a therapeutic strategy for viral diseases. Nat Med 14, 1357–1362 (2008).

46. Moody, M. A. et al. Anti-phospholipid human monoclonal antibodies inhibit CCR5-tropic HIV-1 and induce β-chemokines. Journal of Experimental Medicine 207, 763–776 (2010).

47. Calianese, D. et al. Phosphatidylserine-Targeting Monoclonal Antibodies Exhibit Distinct Biochemical and Cellular Effects on Anti-CD3/CD28–Stimulated T Cell IFN-γ and TNF-α Production. The Journal of Immunology 207, 436–448 (2021).

48. Kenisi, H. et al. Cell surface-expressed phosphatidylserine and annexin A5 open a novel portal of cell entry. Journal of Biological Chemistry 279, (2004).

49. Yang, L. et al. Targeting cancer stem cell pathways for cancer therapy. Signal Transduct Target Ther 5, 8 (2020).

50. Lambrechts, D. et al. A causal relation between bioluminescence and oxygen to quantify the cell niche. PLoS One 9, (2014).

51. Abdeldaim, D. T. & Schindowski, K. Fc-Engineered Therapeutic Antibodies: Recent Advances and Future Directions. Pharmaceutics vol. 15 Preprint at 10.3390/pharmaceutics15102402 (2023).

52. Saxena, A. & Wu, D. Advances in therapeutic Fc engineering - modulation of IgG-associated effector functions and serum half-life. Frontiers in Immunology vol. 7 Preprint at 10.3389/fimmu.2016.00580 (2016).

53. Andersen, J. T. et al. Extending half-life by indirect targeting of the neonatal Fc receptor (FcRn) using a minimal albumin binding domain. Journal of Biological Chemistry 286, (2011).

54. Schatton, T. et al. Identification of cells initiating human melanomas. Nature 451, 345–349 (2008).

55. Hermann, P. C. et al. Distinct populations of cancer stem cells determine tumor growth and metastatic activity in human pancreatic cancer. Cell Stem Cell 1, 313–323 (2007).

56. Galli, R. et al. Isolation and characterization of tumorigenic, stem-like neural precursors from human glioblastoma. Cancer Res 64, 7011–7021 (2004).

57. Al-Hajj, M., Wicha, M. S., Benito-Hernandez, A., Morrison, S. J. & Clarke, M. F. Prospective identification of tumorigenic breast cancer cells. Proceedings of the National Academy of Sciences 100, 3983–3988 (2003).

58. Krishnamachary, B. et al. Hypoxia regulates CD44 and its variant isoforms through HIF-1α in triple negative breast cancer. (2012).

59. Louie, E. et al. Identification of a stem-like cell population by exposing metastatic breast cancer cell lines to repetitive cycles of hypoxia and reoxygenation. Breast Cancer Research 12, 1–14 (2010).

60. Bai, J. et al. HIF-2α regulates CD44 to promote cancer stem cell activation in triple-negative breast cancer via PI3K/AKT/mTOR signaling. World J Stem Cells 12, 87 (2020).

61. Nishikawa, M. et al. Hypoxia-induced phenotypic transition from highly invasive to less invasive tumors in glioma stem-like cells: Significance of CD44 and osteopontin as therapeutic targets in glioblastoma. Transl Oncol 14, 101137 (2021).

62. Pietras, A. et al. Osteopontin-CD44 signaling in the glioma perivascular niche enhances cancer stem cell phenotypes and promotes aggressive tumor growth. Cell Stem Cell 14, 357–369 (2014).

63. Liang, D. et al. The hypoxic microenvironment upgrades stem-like properties of ovarian cancer cells. BMC Cancer 12, 201 (2012).

64. Weng, X., Maxwell-Warburton, S., Hasib, A., Ma, L. & Kang, L. The membrane receptor CD44: novel insights into metabolism. Trends in Endocrinology & Metabolism 33, 318–332 (2022).

65. De Paiva, C. S., Pflugfelder, S. C. & Li, D.-Q. Cell Size Correlates with Phenotype and Proliferative Capacity in Human Corneal Epithelial Cells. Stem Cells 24, 368–375 (2006).

66. Li, Q., Rycaj, K., Chen, X. & Tang, D. G. Cancer stem cells and cell size: A causal link? In Seminars in cancer biology vol. 35 191–199 (Elsevier, 2015).

67. Son, S. et al. Direct observation of mammalian cell growth and size regulation. Nat Methods 9, 910–912 (2012).

68. Li, H., Chen, X., Calhoun-Davis, T., Claypool, K. & Tang, D. G. PC3 Human Prostate Carcinoma Cell Holoclones Contain Self-renewing Tumor-Initiating Cells. Cancer Res 68, 1820–1825 (2008).

69. Schindelin, J. et al. Fiji: an open-source platform for biological-image analysis. Nat Methods 9, 676–682 (2012).

