## Supplemental Fig 1 for "Targeted intracellular delivery of molecular cargo to hypoxic human breast cancer stem cells"

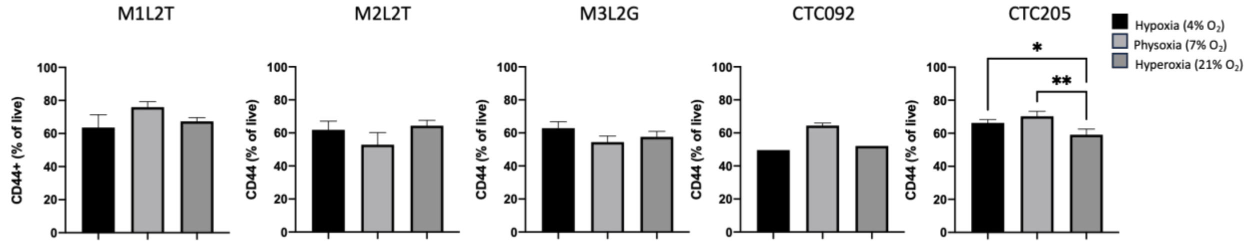


**Supplemental Figure 1. CD44+ populations from each PDX line in different oxygen conditions.** Percent CD44+ cells (from live) determined by flow cytometry for each PDX line used. Cells were maintained in hypoxia (4% $O_2$), physoxia (7% $O_2$) or hyperoxia (21% $O_2$). $*p<.05$, $**p<.01$.
