## Supplemental Fig 2 for "Targeted intracellular delivery of molecular cargo to hypoxic human breast cancer stem cells"

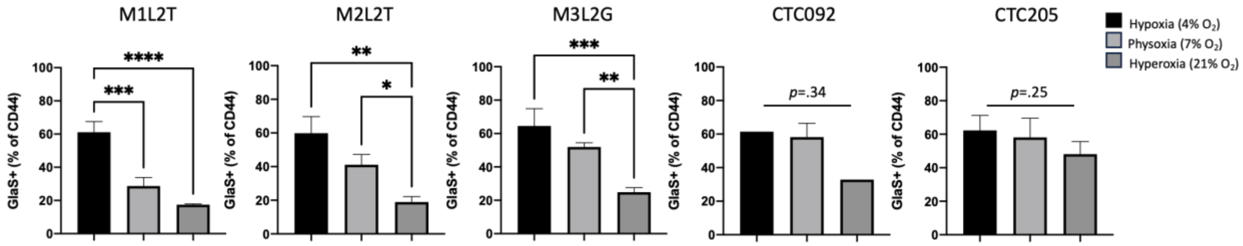


**Supplemental Figure 2. GlaS+ populations from each PDX line in different oxygen conditions.** Percent GlaS+ cells (from CD44+) determined by flow cytometry for each PDX line used. Cells were maintained in hypoxia (4% $O_2$), physoxia (7% $O_2$) or hyperoxia (21% $O_2$). *$p$<.05, **$p$<.01, ***$p$<.001, ****$p$<.0001.
