## Supplemental Fig 3 for "Targeted intracellular delivery of molecular cargo to hypoxic human breast cancer stem cells"

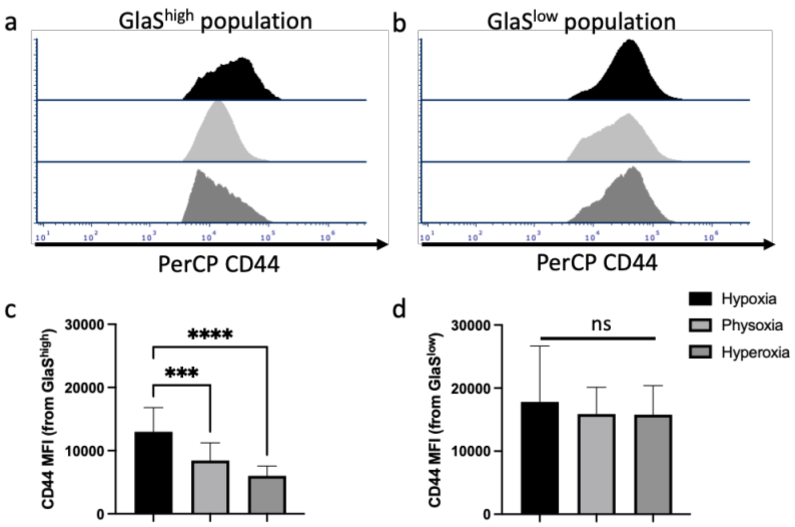


**Supplemental Figure 1. CD44 expression is increased under hypoxia in GlaS$^{high}$small but not GlaS$^{low}$large population.** Flow cytometry histograms demonstrate median fluorescence intensity (MFI) of PerCP CD44 for GlaS$^{high}$small (**a**) and GlaS$^{low}$large (**b**) populations. CD44 MFI was increased in cells cultured in hypoxic vs physoxic and hyperoxic conditions for GlaS$^{high}$small population (**c**). There was no significant difference in CD44 MFI under any oxygen condition for the GlaS$^{low}$large population. n=13 per $O^2$ concentration (n=5 PDX tumor types, n=1-3 technical replicates). ****p*<.001, *****p*<.0001, ns=not significant
