## Supplemental Fig 4 for "Targeted intracellular delivery of molecular cargo to hypoxic human breast cancer stem cells"

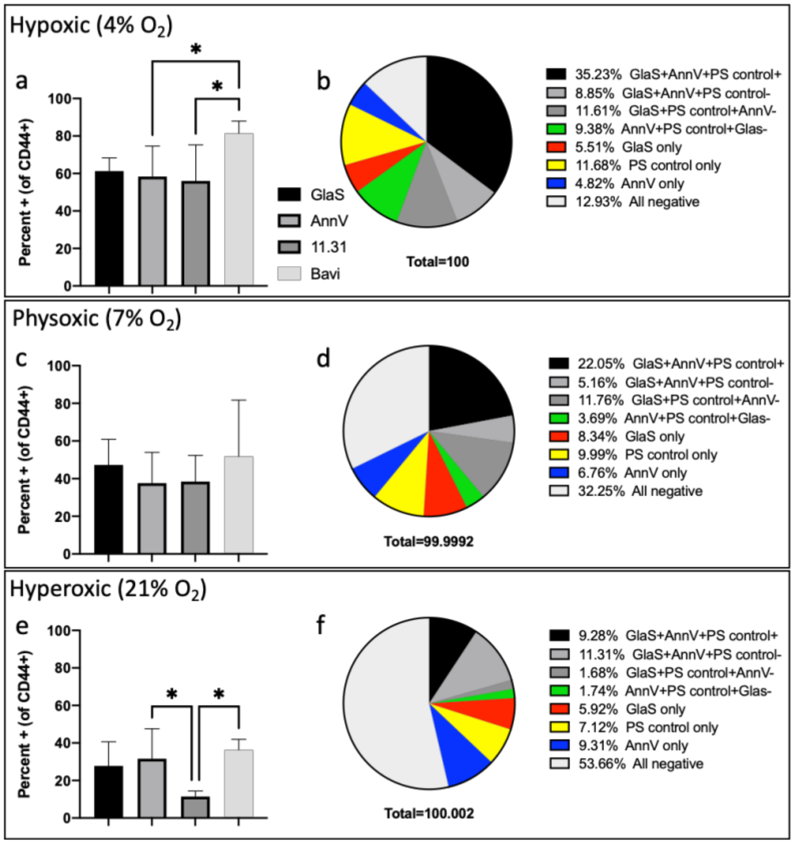


**Supplemental Figure 4 Differences in PS-binding GlaS, Annexin V and PS control Abs.** In hypoxic conditions, there was an increase in CD44+Bavi+ compared to CD44+AnnV+ and CD44+11.31+ cells (**a**). No difference in PS staining was observed for cells cultured under physoxic conditions (**c**). In hyperoxic conditions, CD44+11.31+ cells were decreased compared to CD44+AnnV+ and CD44+Bavi+ cells (**e**). Percent of cells with variations of GlaS, AnnV or PS control Ab staining are demonstrated for hypoxia (**b**), physoxia (**d**) and hyperoxia (**f**). AnnV; n=13 per O$^2$ concentration (n=5 PDX tumor types, n=1-3 technical replicates), Bavi/11.31; n=3-5 per O$^2$ concentration (n=3-5 PDX tumor types, n=1 technical replicate). *$p$<.05
