## Supplemental Fig 5 for "Targeted intracellular delivery of molecular cargo to hypoxic human breast cancer stem cells"

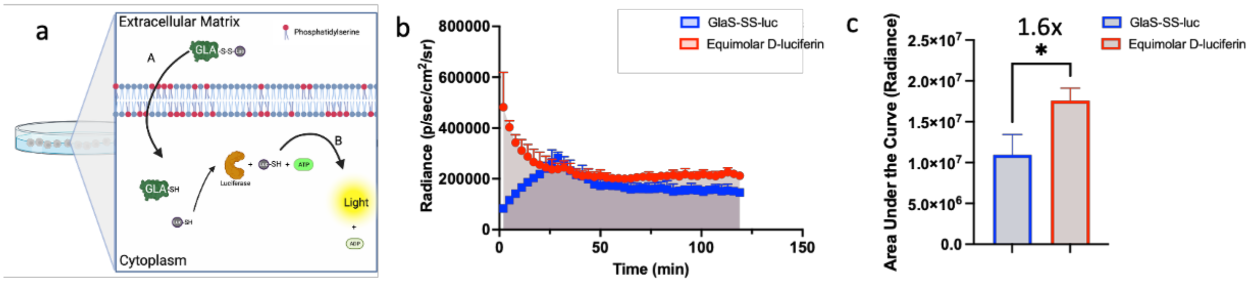


**Supplemental Figure 5. Using bioluminescent imaging as a molecular light switch indicating intracellular delivery of a cargo. (a)** GlaS was conjugated to luciferin via a disulfide linkage (GlaS-SS-luc). Outside of the cell the luciferin remains bound to the GlaS and cannot participate in the bioluminescent reaction. Only once GlaS binds to phosphatidylserine (PS), is internalized, and the disulfide bond is reduced by the intracellular glutathione, the luciferin will be released and there will be light output. **(b)** M1L2T PDX cells were incubated with 1 µg GlaS-SS-luc (blue) or equimolar D-luciferin (red) and imaged over a period of 120 mins. **(c)** Area under the curve (AUC), relating to luciferin delivery and light output (radiance) is compared between GlaS-SS-luc (blue) and equimolar D-luciferin (red). *$p$<.05.
