## Supplemental Fig 6 for "Targeted intracellular delivery of molecular cargo to hypoxic human breast cancer stem cells"

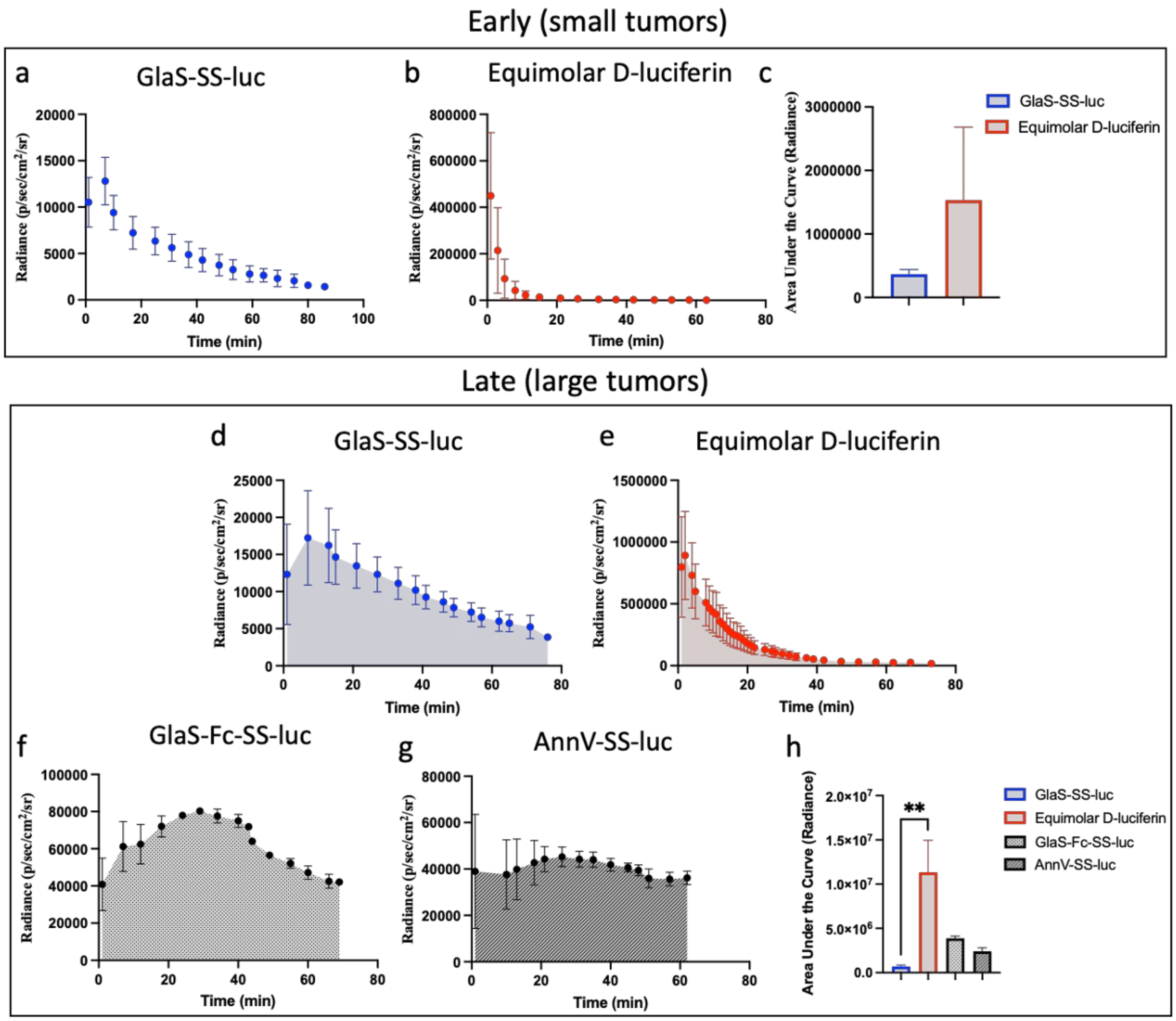


**Supplemental Figure 6. In vivo bioluminescent imaging indicates GlaS-mediated intracellular delivery of a molecular cargo to cancer stem cells.** GlaS-SS-luc or equimolar D-luciferin were administered intravenous into mice bearing small (early) and late (large) M1L2T PDX tumors. In small tumors, there were differences in the kinetics of light output (relating to luciferin delivery) after IV administration of GlaS-SS-luc **(a)** and equimolar D-luciferin **(b).** Area under the curve (AUC) is compared for each material to approximate luciferin delivery **(c).** In large tumors, GlaS-SS-luc **(d)**, equimolar D-luciferin **(e)**, GlaS-Fc-SS-luc **(f)** and AnnV-SS-luc **(g)** were administered IV and bioluminescence was monitored over time. AUC for each material is compared to approximate luciferin delivery **(h)**. **$p$<.01
