## Supplemental Fig 7 for "Targeted intracellular delivery of molecular cargo to hypoxic human breast cancer stem cells"

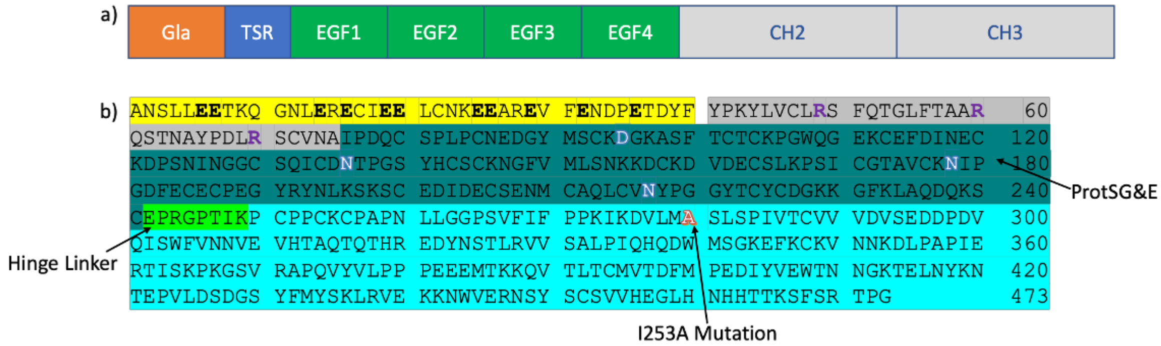


**Supplemental Figure 7. GlaS protein domain structure and amino acid sequence.** GlaS protein was derived from human Protein S and contains the Gla, TSR and four EGF-like domains from Protein S (ProtSG&E) and the murine IgG2A Fc domains CH2 and CH3 **(a).** Amino acid sequence of GlaS: Gla domain (yellow), TSR (grey), EGF domains (green), hinge linker (light green) and Fc domain containing the I253A substitution (light blue) **(b)**.
